# Computer method of biocenotic zoning

**DOI:** 10.1101/2024.11.12.623201

**Authors:** V.V. Sukhanov

**Affiliations:** National Scientific Center of Marine Biology, Far Eastern Branch, Russian Academy of Sciences, Vladivostok, 690041, Russia

**Keywords:** tree stand, nekton, mayflies, regular grid, search radius, species structure, dominant species, correlation coefficient

## Abstract

Algorithm of dividing a territory or a water area into homogeneous areas is described. This area has its own group of communities, similar to each other in terms of species structure. Quantitative statistical indicator, describing quality of such zoning, is proposed. Three examples are given: tree communities in the reserve, nekton in the Sea of Okhotsk and taxocenes of insect larvae in the forest river. Difficulties in the process and interpretation of zoning results, recipes and ideas to solve these problems are discussed.

---

Construction of maps for describing the distribution of areas for many biocenotic complexes, is a compound problem. Because of this, creating even a small number of variants of the same map to choose the best of them involves a large amount of work. In addition, the objective, rigidly algorithmic stages of work (even if they are present) are often accompanied by subjective, not formulated explicitly by the author’s attitudes and assumptions. The quality of such zoning results cannot be verified because there is no objective criterion for this test.

These circumstances served as an incentive for the creation of the algorithm, automating the zoning process based on information about a large number of multidimensional objects, placed in two-dimensional space. The algorithm is illustrated by three examples of zoning: common tree species in the Sikhote-Alin Reserve, species of trawl nekton in the Sea of Okhotsk and insect larvae in 2D space “Distance-Time”.

## Description of the algorithm

The main purpose of zoning is to divide 2D space in fragments containing groups of similar elements. In biocenology such associations are called groupings, biocenotic complexes (Zenkevich, 1963) and taxocenes (when species belong to the same superspecies taxon (Chodorowski, 1959)).

The term “syntaxon” was used as a synonym for these concepts (see definition of Mirkin et al., 1989) in some of our early works on biocenotic zoning. This concept was proposed by the Uppsala school of phytocenologists for the tasks of hierarchical classification of plant communities (the Brown-Blanke system). Linnaeus borrowed the taxonomic system (family - genus - species) was used as a role model. The term “syntaxon” was coined to refer to similar associations of communities in their classifications with a hierarchical structure.

Although spatial classification of communities is the main goal of this work, it does not describe a hierarchical structure. It is not an easy task, similar to the construction of a system of administrative-territorial division of the state territory. The neutral term “group” will be used here. It will represent a homogeneous association, which unites similar biological communities, mostly living in the same area.

The first version of the biocenotic zoning algorithm was created before the era of personal computers and recorded in ancient algorithmic language PL/1 (Sukhanov et al., 1994). The latest versions are realized in the language Free Pascal.

### 1. Creating a Regular Grid

This is the preparatory step, which can be executed separately from subsequent parts of the algorithm. The actual data are descriptions (quantitative collections, samples) of biological communities. They are placed randomly, chaotically in two-dimensional space. We need to streamline this information, turn it into a regular multidimensional data set, presented in the rigid format. One need to divide this space into many identical squares (cells) to do this step. Each of these cells should include information about species abundances located there.

Because we describe a physical two-dimensional space (territory or water area), the shape of the cells should be close to the square to save geometric proportions of future map. It is necessary to line this space with vertical and horizontal lines, where the distance between parallel lines is the same. Thus, many identical square cells are formed, where information about communities will be stored.

The square on the map should be chosen not too big and not too small, so that it can clearly read one character from a pre-selected font. For example, standard sheet of A4 paper, where can be placed a map, should fit several thousand of these square cells.

The space under study is rarely rectangular in shape. It almost always includes one or more closed “blanked” areas. Map cells in these areas should be designated as empty and excluded from calculations. Number *Cell* (see below) designates the number of non-empty cells.

One need to link all the cells to an ordered array of data about the communities in them. This problem is solved by interpolation. The center of the cell is surrounded by a certain search radius *R*. All information in the descriptions (quantitative samples) about species abundances inside this circle, average by weighted arithmetic mean. The statistical “weight” of species abundances is better to choose this way that the “weight” would be smaller if it far from the center of the cell. But it is not necessary.

Search radius *R* may be selected for the following reasons. The more this *R*, the more samples are involved in the calculation of the weighted average in this cell. But the important spatial variations of species abundances can be averaged and eventually leveled. And vice versa, too small radius can lead to absence any information in the cell at all. We often used the formula π*R*^2^ = *C* · *s* · *Cell/Card*, where *Cell* is the number of non-empty map cells, *s* is the area of the cell, *Card* is the total number of samples (actual community descriptions), *C* is a small number of about 5-10. Radius *R* can be set as forced. Then information on species abundances is interpolated into the cell map by searching for all species from the general list.

A simplified version is possible, where the statistical “weight” in the search circle is the same. Then one can use an unweighted average instead of a weighted average.

As shown by personal experience, the creation of a large atlas of maps with a spatial distribution of species abundances is a time-consuming task and it requires sufficient memory and time for computation. Additional difficulties appear for those, who cannot write computer programs. But one can attract well-known cartographic products.

The fast program Surfer-8 can be helpful. It is the old version, but it is a free study program, posted in Internet archives. Free shell QGIS for geographic information systems may be another choice.

Note the following, concluding comments on the first step of the Algorithm. There are two directions in computer visualization - raster and vector graphics (Sukhanov, 2005). Each of them has its advantages and defects.

Zoning map, created by vector graphics, describes many closed areas filled with different shadings and/or colors. Each such area represents the area of a particular group of communities in our case.

The maps presented in this work are done differently, in raster graphics technology. Cells (square boxes, with the help of which our maps are built) are actually the same “pixels” like the variously colored microscopic “grains” on the monitor screen. Only the size of the cells in our case is much larger than the size of these “corns”.

Converting raster images to vector images is a separate task of digitizing the analog images. The description of the methods of its solution is beyond the scope of this article.

### 2. Formation of groups

One need to select a criterion of similarity for a merging of similar descriptions of communities (quantitative samples) in a homogeneous group. This group is assigned its issue (code). This code represents the name of the group. Each map cell receives its own code. It means that all the samples interpolated and averaged in this cell belong to a specific group of communities similar in their structure. In our simplest case, the map cells belong to a group, if their dominant species is the same.

### 3. Definition of reference descriptions for groups

All descriptions with the issue of this group are selected over all cells of the map. According to the above rule, the average “image” of the group is calculated on the basis of the selected set of descriptions. This image is called by “etalon”. It represents typical, the averaged member of this group.

Calculation of the etalon is carried out in this way. Abundance (biomass, number, projective coverage and so on) is summed for each species about all cells, each of which has the issue (name) of this group. The resulting species structure of the etalon is normalized: the sum of all species abundances must be equal to 100%. This procedure for averaging and standardizing the etalon is performed with each of the available groups.

### 4. Elimination of groups

It could be that the group was empty during some iteration of the algorithm because it lost all its members. Another incident could happen if the etalon description of the group ceases to satisfy the criterion, adopted in the step 2. In other words, the etalon is reborn. For example, when the real name of the dominant species no longer coincides with the declared name. Then the ill-conditioned and/or empty group is canceled. And all of its former members are being redistributed to the living, non-liquidated groups. Each such member (a cell of space with a concrete species structure) receives the issue of still living group, which it was closest in its species structure.

It is interesting to observe a similar process of the appearance of reborn concepts in the cultural life of society. The term “ecology” has acquired an overly broad interpretation in recent decades (ecology of intimate relations, ecology of the soul, ecology of native language, ets.).

The meaning of its use is beginning to be lost in biological sciences. The quantitative indicator of the proximity between the etalon and the species structure of the cell is pre-selected by the researcher. We use a correlation coefficient for this purpose (Pianka, 1975). It is the well-known correlation coefficient of Preston, where the correlated arguments are fractions of biomass of the same species, taken as a description of a community in a particular cell, and in the etalon list. This indicator of similarity of species structures of communities is widely known in biocenology (Pesenko, 1982).

### 5. Smoothing the map of similarity with the etalon

We need to create the spatial distribution of the correlation coefficient, the map of similarity of species structures in the cells with the species structure of certain etalon. Atlas of correlation maps is created by running over of all etalons. Then these maps are smoothing out.

Field map smoothing is carried out using a special low-frequency filter. It is a square matrix of dimensions *M*^2^, where *M* is odd number greater than one. This matrix covers a group of *M*^2^ cells of the map at the same time. The elements of the matrix are the statistical weights normalized by the unit. Using these weights, the weighted average of all cells covered by this matrix is calculated. The result is recorded in a cell, combined with its central element. Then the filter moves one step to the side. Its central element is combined with another cell of the unfiltered map and the smoothing operation is repeated. This search continues until each cell of the map is processed by averaging over neighboring cells. The correlation field is smoothed out (maybe repeatedly) and high frequency noise with anomalous emissions are eliminated.

### 6. Renaming map cells

Every cell is again tested after smoothing for compliance with its belonging to one or another still unliquidated group. It is considered that the specific description of the community belongs to the etalon with the greatest similarity. If the past name (code) of the cell does not coincide with the present name after smoothing, the cell is renamed. Then the percentage of renaming cell on the map is calculate. If the number is greater than a predetermined minimum threshold, then we need to go back to step 3 of this algorithm and start the new iteration.

The additional way to interrupt iterations can be provided. The program displays a brief summary of results of the current calculations after the next cycle and asks the question “Continue iterations?”. If the answer is negative, then forced transition to the next step 7 is executed. This way prevents the looping or lack of convergence in the zoning process.

Steps 5 and 6 were combined in later versions of the zoning algorithm. The renaming of cells was carried out immediately after receiving a smoothed map of the correlation field for the next etalon and comparing it with the map for the same etalon, received in the previous iteration. This approach make possible significantly saving computer memory and store maps of correlation fields not for all etalons at the same time, and just for the next one and the last one. Nevertheless, this algorithm remains resource-intensive for real problems.

Let us talk about the optimal number of groups when zoning. The classification also pursues the solution of the practical problem among other purposes: it is the identification of the object’s belonging to a particular group. A good classification allows you to do this quickly (Kafanov, Sukhanov, 1995). Mean time required to identify a randomly selected object is proportional to the log_2_(*N*) with a hierarchically organized classification scheme (dichotomous tree). *N* is total number of groups, contained in the system. Mean searching time is proportional to the value (*N*–1)/2 for non-hierarchical, linearly ordered search system (sequential search). This time is longer than log_2_(*N*) when *N*>6.320.

Thus, linearly ordered classification scheme (English key) is faster with a small number of groups *N*<6-7 than hierarchical system (Swedish key). The boundary *N*=6-7 separating these two keys is close to the number 7±2, known in psychophysiology as the “Miller’s number”.

According to him, usually short-term human memory cannot remember or reproduce more than 7±2 elements. This “magnificent seven” is often found in human practice: seven days a week, seven colors of the rainbow, seven notes in the musical scale, seven wonders of the World, seven major planets in astrology, seven Great Sins, six to seven rounds in a revolver, seven taxonomic ranks (from species to kingdom) in Linnean taxonomy, and at last seven elementary catastrophes of Rene Tom in the mathematical theory of bifurcations.

Miller’s number objectively reflects the upper limit of the capabilities of the human brain by trying to freely and quickly operate with the maximum allowable number of objects or concepts. Non-linear, hierarchically constructed system becomes faster with *N*>7.

Thus, the number of areas depicted on the map for groups of the same rank should not be more than 6-7. This condition should be observed if it is not contrary to other objectives of zoning. Otherwise, the map becomes unreadable. It is much more difficult to work with it and then you need to create a hierarchical zoning system. Selection of the averaging radius *R* or forced termination of iterations can achieve compliance with this condition.

### 7. Map of Group Codes

The final table with an array of all the cells of the map is displayed in an external text file when iterations ended. Each row of the table contains 3 numbers: x and y coordinates of a particular cell and issue (name) of a group characterizing this cell.

Now we need visualize the resulting table as a map. In other words, the results of the zoning must be shown on the graph, partitioning the plane into homogeneous areas. The above-mentioned external mapping program Surfer-8 (Sukhanov, 2005) was involved at this stage in order not to do extra work and get a high-quality map. One can build Classed Post Map (map of symbols, referring to different classes) according to our final table. Each cell on the map is represented by a symbol, codifying a group of biological communities.

In addition, output to the external file displays the final statistical zoning indicators. Among these indicators the most important is the coefficient of similarity (correlations) between the species structure in the cell and the species structure of the etalon characterizing this cell. It characterizes the quality of the zoning result, averaged across all groups: the closer this similarity, the better. The ideal coefficient must be equal to one for correlation index. The Simpson’s species diversity index and other statistics (Pesenko,1982) are also calculated.

## Examples

The zoning maps look almost identical in all the examples. Small differences are in the design of legends to maps. If the table has a fraction of the species less than 0.5%, this number is removed from the table for clarity.

## Sikhote-Alin reserve wood species

This is the first example when development of the zoning algorithm began. Information on the structure of vegetation was collected in 1979 and is stored in the archive of the Sikhote-Alin State Nature Biosphere Reserve (https://sialin.(rf)). The entire territory was divided into 2,596 squares with its area 1.3 km^2^ before computer zoning.

The list of tree species included 28 names. There were 14 most frequently observed species with occurrence at least 0.1%. They were the ones used in the zoning computer program. At the same time, the species were presented initially at a rough taxation level of accuracy. The names of the five species were combined according to the generic name: fir (ayan, korean), larch (okhotsk, komarova), maple (green bark, small leaf, yellow), poplar (scented, maximovicha) and other (10 rarest species). The biomass of these species was expressed in wood units (m^3^/hectare).

It is shown at Fig. 1, how the algorithm reduces the number of cells renaming on the reserve map. The cells belonging to “killed” group are redistributed among the surviving groups after this liquidation. This step leads to a change in their etalon species structures. The change of the etalons in turn changes the relief of their correlation fields. Many cells have to be renamed in the next iteration for this reason, (the curve jumps up a bit at the Fig.1). Then the number of cells renaming drops progressively again, about twice on each iteration. Boundaries between groups begin to “touch” to each other again, until another group of communities is pushed off the map, which brings us back to the upward curve. Iterations end faster when using a larger smoothing filter *M*^2^.

**Fig. 1.**
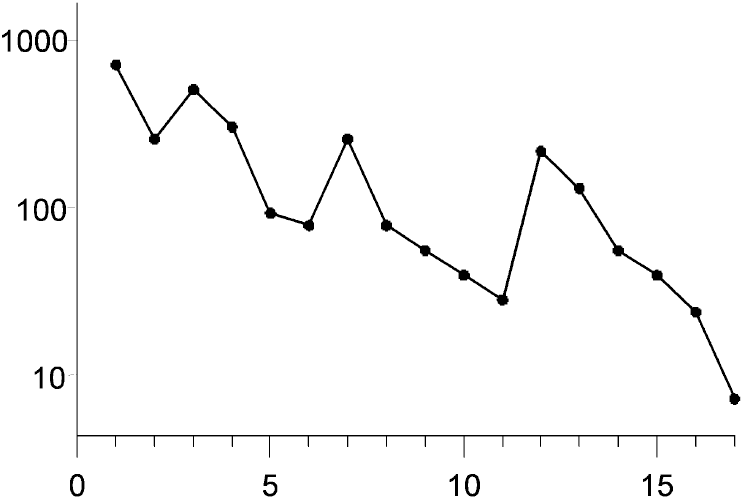
Changes in the number of cell renaming’s of the territory of the Sikhote-Alin reserve (ordinate axis) depending on the number of iterations (abscissa axis)

Species structures of the etalons are given in the Tab. 1, received after completion of the zoning process. There were seven groups in the end. Methodology, used in the classification of phytocenoses (Shennikov, 1964), partly different from the rules, embedded in our algorithm. Nevertheless, these groups can be considered as forest formations.

**Tab. 1.** Species structures of the etalons (% of wood stock) in the Sikhote-Alin Reserve. Only dominants are represented (denoted by bold numbers) in the lists. The little ones are gone. Therefore, the sum of numbers in every column is less than 100%. The areas of reserve occupied by etalons (%) are given before their names

|  |  |  |  |  |  |  |  |
| --- | --- | --- | --- | --- | --- | --- | --- |
| 36.1 cedar | **40.6** | 2.7 | 5.2 | 1.5 | 10.8 | 14.9 | 7.3 |
| 28.3 spruce | 15.1 | **37.3** | 0.8 | 0.1 | 9.1 | 9.7 | 5.1 |
| 14.5 white birch | 4.7 | 3.1 | **36.6** | 12.7 | 15.7 | 7.7 | 7.7 |
| 10.2 oak | 5.1 | 0.7 | 21.8 | **66.0** | 2.3 | 0.5 | 12.4 |
| 5.7 larch | 7.2 | 5.0 | 7.3 | 6.7 | **51.4** | 15.4 | 0.0 |
| 4.3 fir | 10.7 | 2.8 | 2.7 | 0.5 | 6.2 | **30.7** | 3.7 |
| 0.9 stone birch | 2.2 | 5.4 | 1.5 | 0.7 | 0.3 | 1.5 | **55.2** |

Simpson’s diversity of etalons by area is equal to 4.02. It is almost half the total number of etalons. This indicates a noticeable uneven distribution of the reserve area by groups.

The map of tree species zoning is shown at Fig. 2. There are several wide strips along the northeast direction, dedicated the groups. The Sikhote-Alin mountain range extends in the same direction. The small triangular area of the Sea of Japan is presented in the lower right corner of the map.

**Fig. 2.**
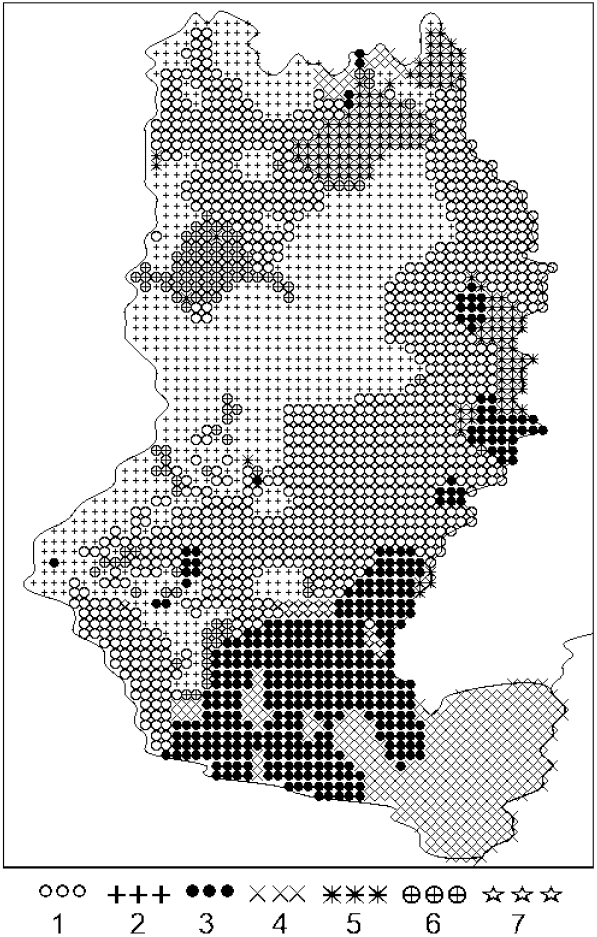
Distribution of tree communities in Sikhote-Alin Reserve. 1 is the cedar, 2 is the spruce, 3 is the white birch, 4 is the oak, 5 is the larch, 6 is the fir, 7 is the stone birch

The increase in the relief height as it moves away from the sea coast causes changes in abiotic factors of external environment. This cause, in turn, leads to regular changes in forest formations. We’ll take the transect northwest and follow it, crossing the Sikhote-Alin range.

Moving from the sea coast, we can see the following changes. The coastal oak up the slope is replaced by white birch and larch, which are replaced then by cedars and fir. After the pass through the Sikhote-Alin range, the height of the relief begins to decline. The fir is replaced by the cedar mixed with silver fir and larch. This picture is partly disturbed by the lateral spurs of the main ridge.

Summarizing this example, it may be noted the following. The level of this map generalization could be made more expressive. That would reduce the number of enclaves and increase the connectivity of areas, although it would certainly be prevented by the high indentation of the relief. In general, the quality of approximation of the forest taxation factual descriptions with the help of seven formations can be considered acceptable. The correlation of species structures in the etalons and in the cells of the map is 0.71. This is a moderate level of similarity between the groups and the actual species structures of the tree stands.

## Nekton of the Sea of Okhotsk

Materials for this example, collected for three decades, owned by the Pacific branch of institute “VNIRO” (Vladivostok, http://www.tinro.vniro.ru/ru/). On their basis it was possible to conduct zoning of nekton biocenoses in the pelagic waters of the Sea of Okhotsk (Sukhanov, Ivanov, 2012). The amount of data involved for this task was 6,727 standard hour trawls.

Biomass of all nekton species, caught in the trawl was determined. The average number of species in the trawl was 6.4, maximum 29 species. Total length of the species list for the Sea of Okhotsk on trawling materials reached 1028 species.

This work on computer zoning was done more than 10 years ago. The taxonomic name of the pollock (the most abundant species in the Sea of Okhotsk) changed during this time. Now it is not *Theragra chalcogramma* but *Gadus chalcogrammus*. Nevertheless, we use its old Latin name. As Shakespeare wrote (Romeo and Juliet), “What’s in a name? (That which we call a rose by any other name would smell as sweet.)”.

The names of the other five most abundant species mentioned below constitute the following list: *Clupea pallasii* is Pacific herring, *Leuroglossus schmidti* is silver fish, *Sardinops melanostictus* is Far Eastern sardine (iwasi), *Mallotus villosus* is capellin, *Oncorhynchus gorbuscha* is hunchback.

Unlike the first example with tree communities of the reserve, the name of the biocenotic complex includes here not only the name of the dominant species, but also the name of a sub-dominant species of nekton. It is a popular approach in biocenology. It was tested on this example with a nekton of the Sea of Okhotsk.

In the first iteration, the program found 170 various community groups, each of which featured a unique combination of dominant and sub-dominant species. After the ninth iteration, there are only seven such groups left. But this simplification did not come free of charge. The average correlation between the species structures of the seven etalons and all the cells of the map decreased from 0.96 to 0.76 (moderate level of similarity). Species structures of the etalons for these final groups are given in the Tab. 2.

**Tab. 2.** Species structures of etalons (% of total biomass) and statistical indicators of groups for the Sea of Okhotsk. The list includes only dominants and subdominants. The sum of numbers in every column is less than 100%. Dash means a fraction less than 0.5%. Groups are coded by abbreviated Latin names. For example, TcCp = *Theragra chalcogramma* + *Clupea pallasii* = pollock + herring.

| Species\Groups | TcCp | TcLs | LsOg | CpTc | TcMv | SmLs | MvTc |
| --- | --- | --- | --- | --- | --- | --- | --- |
| *Theragra chalcogramma* | 45.4 | 75.7 | 11.4 | 20.2 | 47.5 | 2.6 | 10.8 |
| *Clupea pallasii* | 16.8 | 4.0 | 2.2 | 62.5 | 7.1 | 1.3 | 8.3 |
| *Leuroglossus schmidtii* | 12.8 | 6.1 | 27.5 | - | - | 14.3 | - |
| *Sardinops melanostictus* | - | - | 4.6 | - | - | 39.5 | 0.6 |
| *Mallotus villosus* | 1.2 | 1.0 | - | 3.8 | 23.1 | - | 54.5 |
| *Oncorhynchus gorbuscha* | 3.8 | 1.7 | 17.7 | 0.9 | - | - | 5.8 |
| Statistics\Groups | TcCp | TcLs | LsOg | CpTc | TcMv | SmLs | MvTc |
| Catch ton/hr | 10.5 | 23.0 | 7.1 | 20.2 | 7.0 | 10.7 | 9.9 |
| Species diversity in catches | 3.3 | 2.2 | 4.5 | 2.4 | 3.0 | 3.1 | 2.9 |
| Correlation of etalon with cells | 0.81 | 0.84 | 0.69 | 0.71 | 0.71 | 0.59 | 0.57 |
| Species diversity of etalon | 4.4 | 1.8 | 7.7 | 2.5 | 3.7 | 5.5 | 3.4 |
| Water area % | 43.0 | 18.8 | 13.3 | 12.9 | 5.3 | 4.1 | 2.6 |

Analysis of group area map (Fig. 3) indicate a large number of enclaves. The habitats of all the groups were torn apart. We have to explain, why did this happen.

**Fig. 3.**
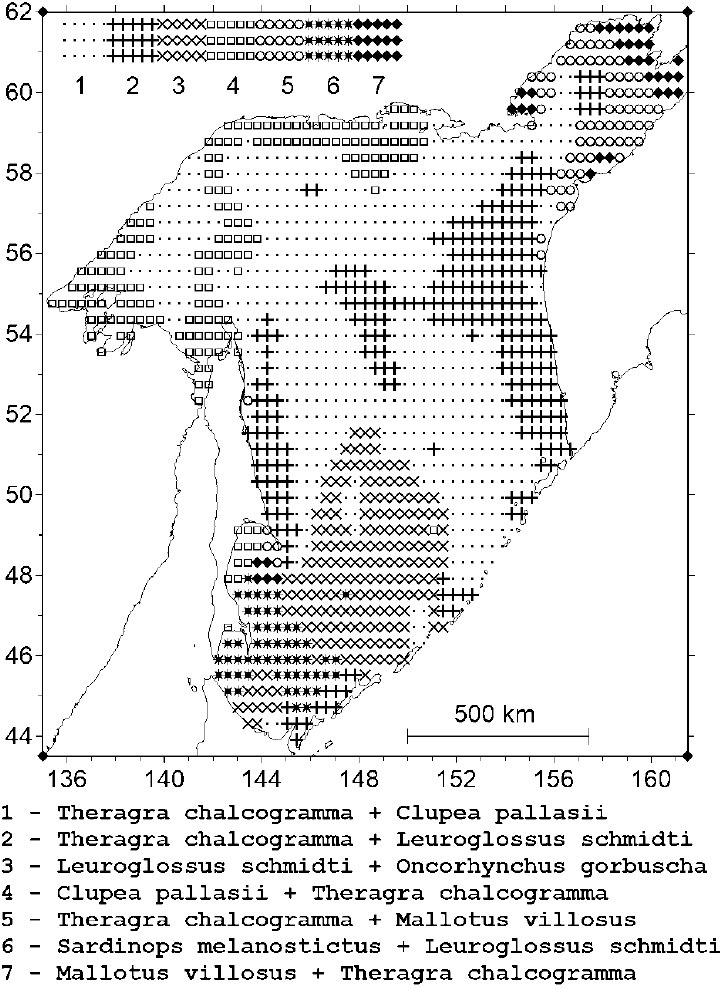
Distribution of trawl nekton communities in the Sea of Okhotsk. The share of the water area and its number are indicated before the name of the group. The name includes dominant and sub-dominant species

The Sea of Okhotsk is a three-dimensional object, not only horizontally along geographical coordinates, but also in depth. Mean depth is 821 m there, maximum 3916 m.

Nekton species don’t just live in epipelagic zone, but also at great depths. Fig. 4 shows the ecological niches of the seven most common species, their total biomass is on average 97% of the total biomass of nekton. These niches were built as follows.

**Fig. 4.**
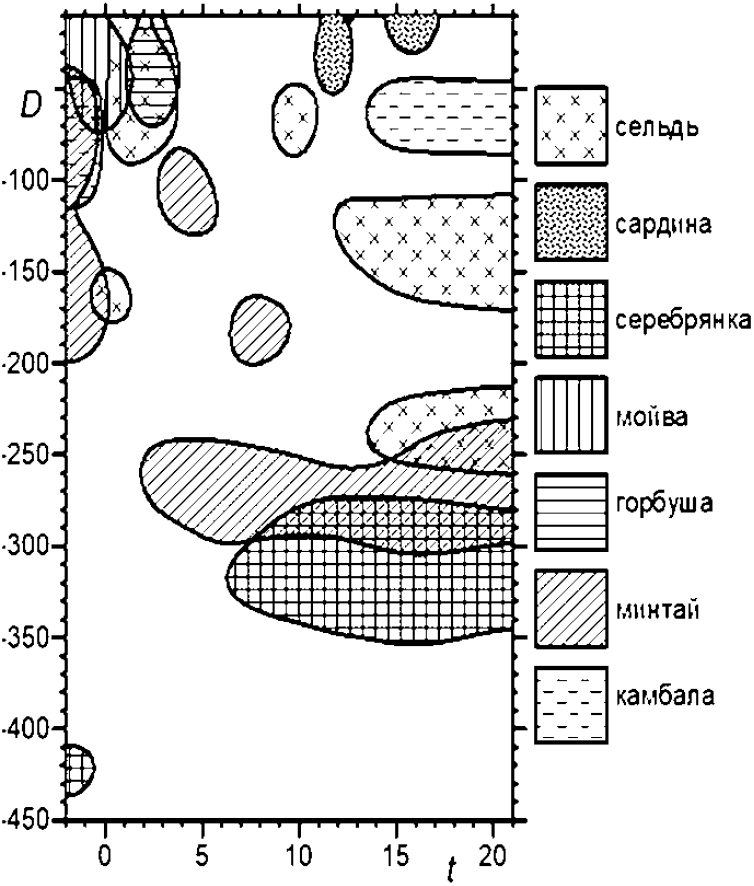
Ecological niches of common species of nekton in the Sea of Okhotsk in the space of two environmental factors. Vertical axis *D* is the depth of detection (m), horizontal axis *t* is the water surface temperature (°C)

Formalization of the term ecological niche was proposed by Hutchinson (1957). It is based on the concept of a function of a set of arguments. These arguments describe environmental factors that are vital to the species. The function itself is the “density of fitness”(Pianka, 1981), that is physiological or population-demographic reaction of a species to concrete values of these factors. Maximum of this density indicates the center or optimum of the niche. As we move away from this optimum, the density of fitness decreases to the limits of ecological niche of the species.

Biomass of the species in the catch was selected as the fitness density. Two environmental factors were always measured at each trawl. It is the depth of the trawling and the temperature of surface water. The first factor reflects the habitat of the species on the vertical axis of depth. The second factor serves as an indirect marker of seasonal changes in the abiotic environment: the temperature is low in the winter, high in the summer and takes intermediate values in the demi-season periods (spring or autumn).

The choice of surface temperature (abscissa axis) is not entirely successful: the temperature is low and almost constant at a great depth. It plays the role of seasonal time here. This is a cyclic variable (from January 1 to December 31 and then January 1 again), and you need a polar coordinate system to represent it. Although it is possible combine a depth and a circle with cyclic seasonal time (using the Wind Rose), it is unusual. Therefore, described option was chosen here.

The boundaries of species niches were defined as follows. A graph of the distribution of its biomass was constructed in the coordinates “surface temperature - depth” for each species. A point with maximum biomass was found on the graph, and it was assigned the optimum Hutchinson’s niche. A third of this amount (33% of the maximum) was conditionally accepted as boundary of the ecological niche. The isoline running along this boundary was shown in the graph. The entire inner region of the species econiche was filled with a special type of shading. Then the niche graphs for the 7 common species were combined into Fig. 4.

It would be possible to discuss in detail the regularities of the mutual arrangement of these econiches. But we need only one fact here. There is a strong overlap of niches in seasonal aspect at any time of the year. It means that projections of niches on the plane of geographical coordinates almost always lead to the imposition of species ranges on each other that is on the map of zoning (Fig. 3). For example, the sardine, flounder, herring, pollock and silver fish can be included in the names of the same groups in various pairs in the warm season, although not all of them live together. This results in a pseudo-random mishmash of grouped ranges in the plane.

There are several ways out of this difficult situation. First, you can do layered zoning, when each map layer is assigned to its own depth layer. Second, one can make a series of maps for different intervals of seasonal time. Third, one can use different colors to represent group habitats at different depths. In general, the solution of these problems with the help of software will require considerable additional efforts and not the fact, that the result will be successful.

Fourth, one can zoned in three-dimensional space “longitude — latitude — depth”. The program Voxler-4 may be useful for this purpose (Sukhanov, Kiyashko, 2021). We had to come across difficult new and still unresolved problems of zoning in 3D space in the process of solving the challenge. The main problem is that the methods for constructing a three-dimensional lattice are still poorly developed. This space must be filled with a 3D parallelogram (bricks) and not with a 2D squares.

In addition, there are difficulties of visualizing objects in three-dimensional space. Conversion of 2D in 3D space is not simple. Our eyes are adapted for processing not three-dimensional, and generally two-dimensional information. The retina is 2D organ, on which real 3D images are projected. Eye accommodation allows you to determine the distance to objects in a three-dimensional environment. But this property is useless when building volumetric models on a computer: the monitor screen is two-dimensional and has no real depth. We need to use spatial imagination. But this is not a job for the eyes and for the brain. It’s not easy.

Another way to solve this problem is to use a simpler zoning scheme, as in the first example of a forest reserve. The name of the group can be determined by the names of only the dominants. We got the following in the end.

29 groups were identified in the first iteration with a correlation coefficient of the etalons with the actual descriptions, is equal to 0.92. After the sixth iteration, seven groups remained alive with a correlation coefficient of 0.76. At the same time, the two smallest groups occupied only 2% of the water area. They were forcibly removed and their cells were redistributed to five larger surviving groups. The correlation coefficient did not decrease after that and was still equal to 0.76. The specific structures of the five etalons and statistical indicators are given in the Tab. 3.

**Tab. 3.** The species structure of etalons (% of total biomass) and statistical indicators of the groups for the Sea of Okhotsk. Only dominants are represented in the lists, denoted by bold numbers. The sum of numbers in every column is less than 100%. Dash means a fraction less than 0.5%.

| **Species\Groups** | **Tc** | **Ls** | **Cp** | **Sm** | **Mv** |
| --- | --- | --- | --- | --- | --- |
| *Theragra chalcogramma* | **65.5** | 19.9 | 26.4 | 9.5 | 22.5 |
| *Leuroglossus schmidtii* | 4.0 | **26.3** | 0.6 | 17.6 | - |
| *Clupea pallasii* | 12.7 | 3.5 | **56.0** | - | 7.6 |
| *Sardinops melanostictus* | - | 0.6 | - | **39.7** | - |
| *Mallotus villosus* | 2.5 | 0.3 | 3.5 | - | **42.3** |
| **Statistics\Groups** | **Tc** | **Ls** | **Cp** | **Sm** | **Mv** |
| Catch ton/hr | 18.4 | 5.0 | 18.3 | 15.6 | 8.5 |
| Species diversity in catches | 2.2 | 4.6 | 2.3 | 3.3 | 3.2 |
| Correlation of etalon with cells | 0.84 | 0.72 | 0.74 | 0.59 | 0.61 |
| Species diversity of etalon | 2.2 | 7.4 | 2.6 | 4.8 | 4.1 |
| Water area % | 39.1 | 31.3 | 17.7 | 7.2 | 4.7 |

This variant of zoning shows Fig. 5. The number of enclaves has decreased, areas began to look easier without degrading the quality of the material approximation (the correlation coefficient was the same in the first and in the second case). Apparently, that zoning with dominant species only was easier to perceive.

**Fig. 5.**
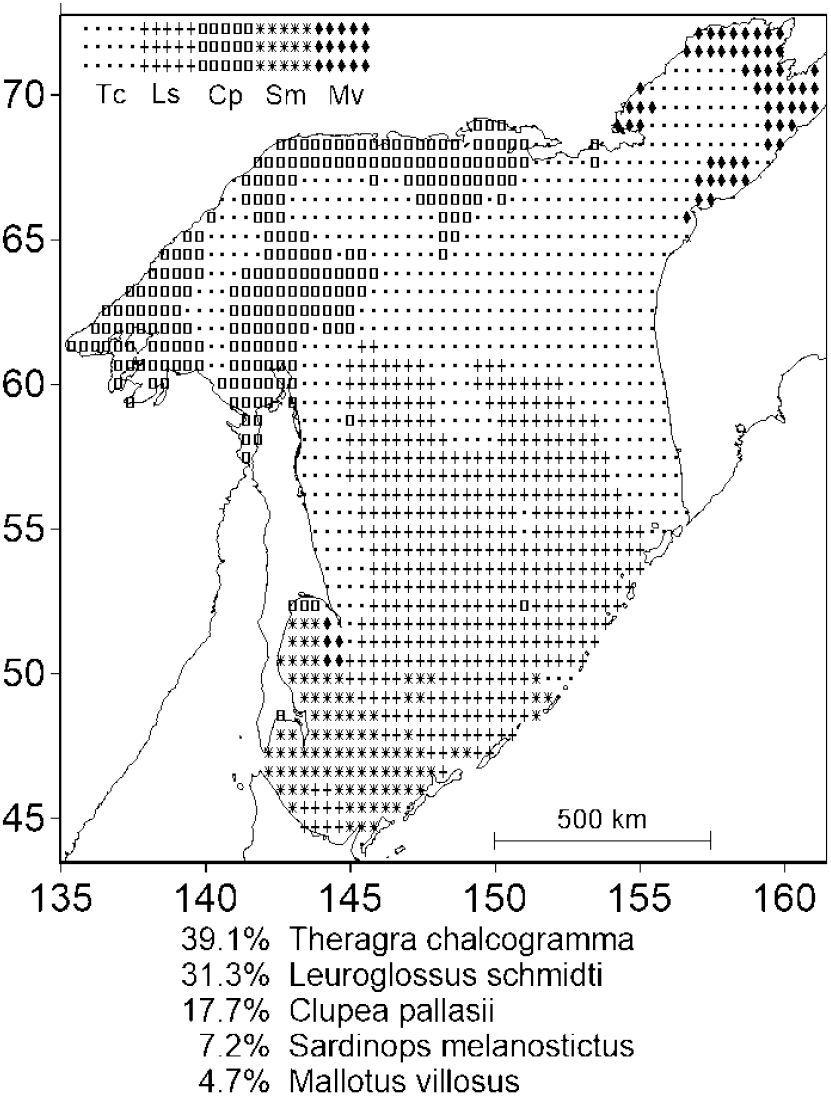
Distribution of nekton communities in the Sea of Okhotsk. The proportion of the water area is indicated before the name of the group. The name includes only the dominant species

## Mayfly larvae in a forest river

The third example focuses on the larva communities (*Insecta: Ephemeroptera*), inhabiting the bottom of the shallow fordable forest river Komarovka on the south-western slope of Sikhote-Alin (Primorsky Krai, swimming pool of the river Razdolnaya). The material was collected from May to October using samplers at 11 stations, installed on the longitudinal profile of the watercourse from the source to the middle course of the river. This section is at the top of the river basin, located within the Ussuri reserve. It is entirely within the security zone, without human impact. The list includes 70 species founded in 44 samples.

We have two factors of the external environment: seasonal time and distance from the source of the river. These environmental factors have different units of measurement (distance is measured in kilometers, and time in months). Therefore, the direct application of the zoning algorithm to this 2D space is impossible. Although there are examples of successful work with such spaces directly: “Soldiers, we are digging now the trench from the fence to the lunch”.

First, we got rid of the units to solve this problem. The average time was subtracted, and then the result is divided by the standard deviation in time for all the observed moments of time. The result was a dimensionless quantity with zero mean and a single standard deviation.

We did the same with the distance from the source of the river. We took logarithms of distance before this procedure. This distortion of the spatial coordinate was necessary because the sharpest and most noticeable changes in the species structures of communities are observed precisely in the upper reaches of the river. We wanted to study them more closely, and logarithms stretch small distances and compress large distances.

When the zoning was completed, we’re back to the original coordinates of space and time, using reverse transformations. They are presented on the final map.

Cell renaming stopped after 17 iterations. The number of final groups turned out to be equal from the original ten to six (Tab. 4). Total correlation between the etalons and the cells of the map was equal to 0.87. That is not a bad indicator.

**Tab. 4.** The species structure of etalons (% of biomass) and statistical indicators of the groups for mayfly larvae in the taiga river. Dominants are marked in bold. Secondary species omitted; dash means a fraction less than 0.5%. The sum of numbers in every column is less than 100%.

| Species\Groups | Da | Bs | Dc | Eo | Ep | Ea |
| --- | --- | --- | --- | --- | --- | --- |
| *Drunella aculea* | **30.7** | 2.8 | 17.8 | 13.1 | 11.8 | - |
| *Baetis sibiricum* | 4.7 | **39.2** | - | - | 0.2 | - |
| *Drunella cryptomeria* | 4.8 | 3.9 | **30.4** | 17.9 | 23.0 | - |
| *Ephemera orientalis* | 1.1 | - | 6.5 | **19.8** | 0.2 | - |
| *Epeorus pellucidus* | 1.3 | 0.3 | 11.0 | 6.3 | **30.5** | 3.2 |
| *Ecdyonurus aurarius* | 0.5 | 6.1 | - | - | - | **40.3** |
| Species diversity in samples | 6.7 | 5.4 | 6.0 | 7.6 | 5.2 | 3.5 |
| Correlation of etalon with cells | 0.80 | 0.86 | 0.94 | 0.91 | 0.89 | 0.86 |
| Species diversity of etalon | 8.9 | 5.4 | 6.6 | 8.7 | 6.1 | 3.6 |
| Area % | 29.4 | 20.4 | 16.9 | 16.2 | 14.4 | 2.6 |

The correlation of the species structures with the cells on the map is quite large and varies little from group to group. The greatest diversity of etalons was in the largest complex *Drunella aculea* (8.9) and the smallest in the little *Ecdyonurus aurarius* (3.6), This is no surprise. The same conclusion can be drawn about species diversity in the map cells.

Fig. 6 shows the final map of biocenotic complexes. Coherence of occupied groups of areas can be considered high. Only two groups of six (*Drunella aculea* and *Epeorus pellucidus*) have small enclaves in neighboring areas.

**Fig. 6.**
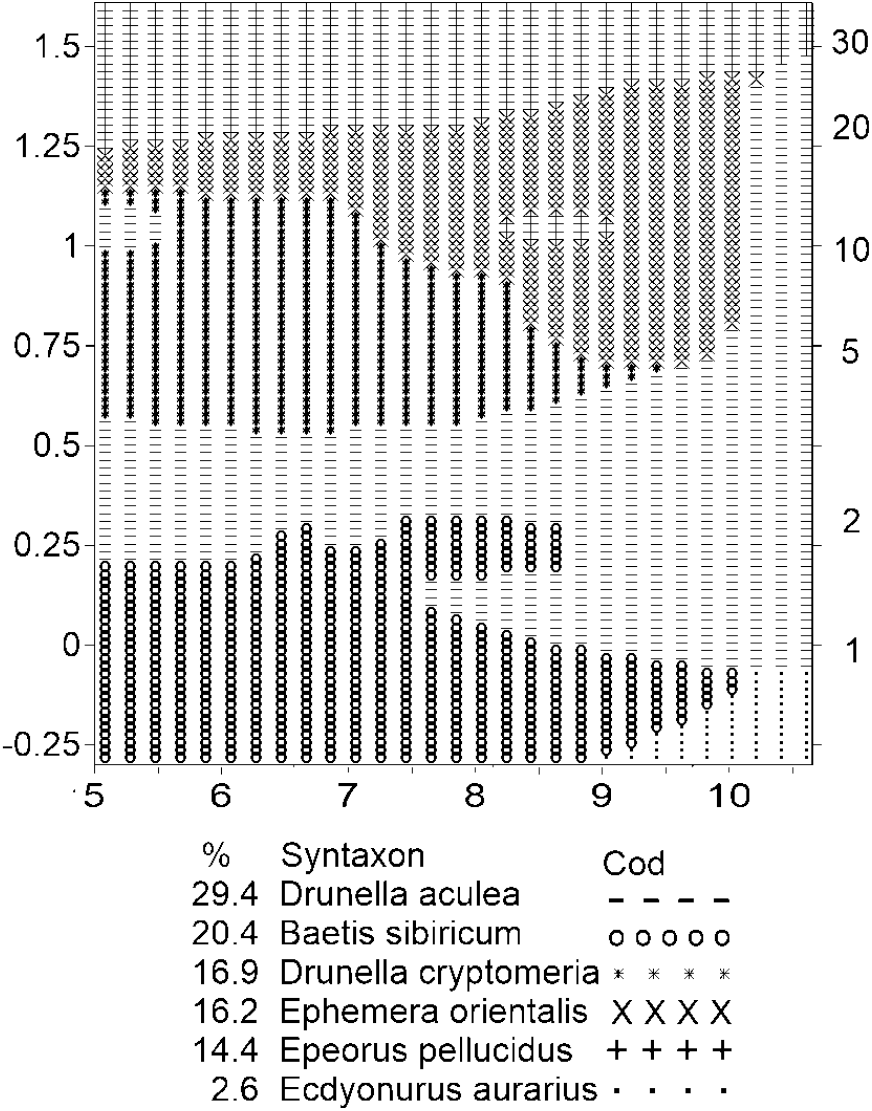
Distribution of mayfly larvae populations in Komarov River. The share of the area in %, then the name and the group symbol on the map are indicated. Abscissa axis is the seasonal time (month), ordinate axis on the right is the distance from the source (km), the ordinate axis on the left is the same distance in decimal logarithms

The habitats of most groups are observed in almost the entire time range from May to October. Just a small complex *Ecdyonurus aurarius* was registered in September-October near source of river.

The main changes in the areas do not occur in seasonal time but in the space. This is clearly visible on the map: groups of horizontal bands replace each other in the process of moving along the river. This is not surprising: the larvae of mayflies almost do not migrate.

The all-season group *Epeorus pellucidus* take up position in the range from about 30 to 20 km from the source. Complex *Ephemera orientalis* is also all-season but it reaches its greatest development from August to mid-October. Group *Drunella cryptomeria* takes up position next to the source from Spring to September, it is common in the area of about 15 to 3 km. Complex *Drunella aculea* observed from spring to early October in the range of 2-3 km from the source. It spreads along the river almost everywhere, thereby disrupting the pattern of successive shifts of groups along the river. Group *Baetis sibiricum* localized directly at the beginning of the river and ends at a distance of about 2 km, it is observed from spring to autumn. Small group by area *Ecdyonurus aurarius* is founded in October near the source.

Another zoning option was calculated where the name of the complex is not given by one, and the names of two species at once: dominant and subdominant. The initial number of groups before the zoning process began was 44, i.e. each sample was identified as a unique group. After completion of the calculations, their number was reduced to 7. Let us give the final list of group names for the option “dominant + subdominant”.

The volume of the group is indicated in parentheses, as a percentage of occupied cells:

*Drunella cryptomeria* + *Drunella aculea* (23.4),

*Baetis sibiricum* + *Ameletus cedrensis* (20.6),

*Drunella aculea* + *Cinygmula spp*. (19.1),

*Drunella aculea* + *Cinygmula supporensis* (15.2),

*Drunella cryptomeria* + *Epeorus pellucidus* (7.9),

*Epeorus pellucidus* + *Drunella cryptomeria* (7.2),

*Baetis sibiricum* + *Epeorus (Iron) maculatus* (6.6).

The connectivity of habitats in biocenotic complexes was low here. Only two groups had intact habitats, the other five groups had enclaves in neighboring areas. Nevertheless, the average correlation of the etalons with the species structures of the samples was 0.89. That is a bit more than in the first zoning, where the names of the groups were given only by the names of dominant species. Thus, the etalons also approximated the species structures in the space cells quite well here.

Nevertheless, it was not appropriate to give a map of complexes and a table of species structures for etalons in this version. The map looked convoluted and showed no obvious spatial patterns. The table of species structures for the etalons also turned out to be illustrative example. The zoning option with double names of the dominant + subdominant groups turned out to be unsuccessful in our opinion. This failure was not due to the fact that the larvae lived in three-dimensional space, as in the example of a certain Sea of Okhotsk: they live at the bottom of the river.

The quality of approximation of species structures in the first variant of zoning turned out to be only 2% worse, than in the second. But understandable and easily interpreted map was in the first version. In the second version with the addition of sub-dominants, an intricate hodgepodge of ranges came out like a rustic scrappy blanket. Excessive complexity of the task has led to a deterioration in the visibility of the results.

Based on all three examples, an impression can be created that zoning only on dominant species should always lead to better results compared to the option of “dominant + subdominant”. But it may be that the distribution of species abundance deviates greatly from the typical geometric progression. In other words, let the dominant species is the same in almost all samples. Then zoning only by dominants would be pointless, and you have to attract sub-dominant species.

## Conclusion

The paper discusses the results of 30 years work with the computer method, enabling biocenotic zoning to be performed. Probably this is the first algorithm solving such problems in geography. It can be used not only in ecology but also in socio-economic, medical, meteorological services, geological and other sciences, where it is necessary to classify space into regions with homogeneous content. The algorithm is implemented in the original program, but in the non-commercial version: there is no user-friendly interface here.

Experience with the algorithm has shown that the best results of zoning are obtained when group names are given by dominant species, without sub-dominant species. Then the maps are simpler and clearer, group habitats are more connected and with fewer enclaves. Thus, single-level zoning of biocenotic complexes (by dominants) so far seems to be more reliable than hierarchical. Nevertheless, in special cases, meaningful results can be obtained only with the involvement of the option “dominant + subdominant”.

Algorithm successfully works in unusual 2D spaces where the coordinates on the maps can be not only geographical, they may have different units of measurement. It also allows you to work with non-quantitative species lists, when the species have no information about their abundance, and there are only facts about their presence or absence at concrete points of material collection. One can use here the unit frequency of occurrence of a species at a given point instead of the index of species abundance, and then it is smeared across the space with a smoothing filter.

An objective criterion of the quality of the received zoning is proposed. This is the correlation coefficient between the species structures of real communities, represented in the map cells and between the etalons - the average species structures of the detected groups. This ratio provides an objective indicator describing the quality of zoning in the region. Ideally it should be equal to 1. In addition, we can consider the fact, that for the correlation coefficient, in principle it is possible to calculate the statistical error. The downside is that it is not clear, how to estimate the real number of freedom degrees after manipulation with smoothing and interpolation. Therefore, the testing of statistical hypotheses will have to be postponed until this problem is resolved.

The mean correlation coefficient turns out to be close to the estimate of 0.7 – 0.8 as a criterion for the quality of zoning in our examples. If we square this, then the determination factor results roughly. It shows how much of the empirical variance is due to zoning, not caused by random noise. Determination coefficient at about 0.5 – 0.6 is slender. What happened is what happened.

In the process of working on the zoning algorithm, we encountered new and still unresolved problems of zoning in three-dimensional space. There are difficulties (including psychological) in the projection of real three-dimensional objects on the plane of the monitor. This is a separate circle of mathematical problems (Rauschenbach, 1986).

But the main difficulty consists of still poorly developed calculation methods for constructing a three-dimensional grid (lattice), filled with aThis work was funded from the budget of the Institut e“bricks”, 3D parallelograms (voxels) but not square cells (large pixels).

## Gratitude

The author thanks Dr. O.A. Ivanov (Tinro) for actual material marching the Fig. 4, and also Ph.D. T.C. Vshivkova (FNCB FVO RAS) for data on the mayfly larvae. Thanks to the leading specialist of VNIRO P.O. Emelin for critical review of the manuscript.

## Funding of work

This work was funded from the budget of the Institute. No additional grants have been received to conduct or direct this particular study.

## Compliance with ethical standards

There are no studies of humans or animals.

